# CRISPR-Cas9 and PiggyBac Mediated Genetic Modification of Sand Fly Vectors Targeting Olfactory and Non-Lethal Phenotypic Genes

**DOI:** 10.64898/2026.02.24.707685

**Authors:** Rhodri T. M. Edwards, Luke Brandner Garrod, Tapan Bhattacharyya, Barbora Vomáčková Kykalová, Erich Loza Telleria, Matthew E. Rogers, Thomas Walker, Petr Volf, Matthew Yeo

**Author notes:** To whom correspondence should be addressed: Matthew Yeo, Ph.D., Department of Infection Biology, London School of Hygiene and Tropical Medicine, Keppel Street, London WC1E 7HT, UK, Ph.

## Abstract

*Leishmania* spp. are protozoan parasites transmitted by female sand flies (Diptera) that cause a spectrum of devastating human pathologies affecting millions. Control of leishmaniases has proved immensely difficult: vector control strategies remain challenging and available chemotherapeutics imperfect. Gene editing of insect vectors offers prospects to interrupt disease transmission; for example by introducing antiparasite effectors or heritable modifications of genes implicated in fecundity. Here we present convergent evidence for successful gene editing using both CRISPR–Cas9 and PiggyBac approaches in two medically important sand fly species (*Lutzomyia longipalpis* and *Phlebotomus papatasi*). Targeted mutagenesis in G_0_ and G_1_ generations is supported by three independent lines of evidence, namely observable phenotypic changes, PCR-based detection assays, and confirmatory *in silico* algorithmic analysis of gene sequences which together provide a robust demonstration of targeted genome editing.

## Introduction

CRISPR has transformed gene editing in insects of medical and veterinary importance due to a highly targeted and relatively simple application in comparison to previous methods (Dong et al. 2015). An important utility of CRISPR-Cas9 are the approaches to interrupt disease transmission. CRISPR-Cas9 based gene-drives that enable rapid spread of introduced heritable traits in fast-reproducing insect populations is a promising approach (Bier 2022). Two favoured applications of gene-drives are i) targeting genes that affect fecundity to suppress populations below the threshold required for parasite transmission (Hammond, Pollegioni, et al. 2021; Hammond et al. 2016), and ii) promoting population modification through the expression of introduced anti-parasite effectors that confer resistance to associated pathogens (Adolfi et al. 2020). Both approaches show great promise in laboratory and large-cage settings.

Despite rapid progress in gene editing methods in medically important mosquito vectors, including *Aedes* sp. (Anderson et al. 2023)., *Anopheles* sp. (D’Amato et al. 2024)., and *Culex* sp. (Harvey-Samuel et al. 2023), advances in other medically important insect vectors remain limited, including sand flies that transmit the protozoan parasite *Leishmania*. Presently, control of sand flies, including *Lutzomyia longipalpis* and *Phlebotomus papatasi*, remains challenging (Kumari et al. 2025) due in part to their highly dispersed, low density terrestrial breeding sites (Balaska et al. 2021). Chemotherapeutics to treat leishmaniasis are imperfect and often not available to those in need (eBioMedicine 2023). Therefore, demonstration of gene editing of endogenous genes and the insertion of exogenous genetic cargo is an important step toward advancing sand fly biology and exploring new opportunities to control leishmaniasis, including gene drive development.

Currently there is a relative dearth of published research on the application of CRISPR-Cas9 in sand fly vectors. Optimisation of embryonic microinjection techniques for CRISPR-Cas9 editing has been described (Martin-Martin et al. 2018; Jeffries et al. 2018; Louradour et al. 2020). CRISPR-Cas9 mediated transgenesis has been demonstrated in a single landmark study targeting a *P. papatasi* transcription factor (NF-κB) and the Relish gene (*Rel*), a component of the immune deficiency (IMD) pathway (Louradour et al. 2019). In that report, recombinant Cas9 protein was directly injected together with sgRNAs derived by *in vitro* transcription to achieve targeted knockouts. The resulting *Rel* knockout colonies were difficult to maintain, and the number of injected embryo survivors was low, although efficiency of transgenesis was relatively high. To date, there are no published reports of successful genetic modification in *L. longipalpis*, the primary vector of visceral leishmaniasis in the Americas, using either CRISPR-mediated transgenesis or PiggyBac transformation.

Here we demonstrate the microinjection of CRISPR plasmid constructs and assess subsequent gene edits in two major sand fly vectors, providing the first demonstration of targeted CRISPR-induced mutagenesis in *L. longipalpis*, resulting in phenotypic mutation, as well as mutagenesis of olfactory genes involved in host seeking. Additionally, we report PiggyBac-mediated cargo insertion with subsequent inheritance in both *L. longipalpis* and *P. papatasi* following large-scale embryo microinjection. We provide convergent lines of evidence for successful mutagenesis in emergent transfected sand flies, namely targeted phenotypic changes, PCR-based detection assays, and confirmatory *in silico* algorithmic analyses of sequence data. Together, these findings provide robust support for the feasibility of gene editing in sand flies.

## Methods

### Identification of gene targets to affect

Candidate genes were chosen based on the likelihood of a phenotypic effect and non-lethality upon knockout. Genes with conserved function across multiple insect orders were selected using FlyBase (flybase.org) and VectorBase (vectorbase.org), and orthologues within *L. longipalpis* and *P. papatasi* identified. Orthologues were assessed via Clustal Omega (ver. 1.2. 3.) protein alignment against model organisms to infer gene ontology searching on AmiGO 2 (ver. 2. 5. 17) using template genomes lonJ1.6. and papI1.6. hosted on VectorBase. Briefly, candidate target genes likely to elicit a non-lethal phenotypic change were *ebony* (*e*), *vestigial (v), and rudimentary (r)*. Separately candidate olfactory genes implicated in host seeking were chosen including *Orco, Gr2* and *Ir8a* (Supplementary Table S1).

### Identification of gRNAS

ChopChop (Labun et al. 2021) was used to identify gRNA sequences using standard settings, which were selected based on efficiency scores (Doench et al. 2014). Functional gRNAs were assessed by an *in vitro* cleavage assay confirming activity. gRNAs were transcribed using the Engen Transcription kit (NEB, USA) following the manufacturer’s protocols. Synthesised sgRNAs ability to cleave was assessed against amplified template DNA. Successful cleavage evidenced by digestion of template DNA was visualised by gel electrophoresis (Supplementary Figure S2).

### CRISPR and piggyBac plasmids

For CRISPR induced knockouts the pDCC6 vector served as the backbone. The plasmid carries a gRNA and Cas9 binding scaffold expressed by a *Drosophila* U6 pol III promoter for RNA transcription and a human-codon-optimized Cas9 coding sequence (hCas9) promoted by an hsp70 pol II promoter (Gokcezade et al. 2014). pDCC6 was modified to produce a panel of plasmids targeting different genes (Supplementary Table S3). The dU6-2 (FBgn0266758) promoter was replaced with an endogenous U6 promoter (*L. longipalpis* and *P. papatasi*) via Gibson Assembly following manufacturer’s instructions (Gibson Assembly® Cloning Kit, NEB, USA). The region 400bp upstream of the U6 start codon encompassed key promoter sequences including the pol III proximal sequence element A (PSEA) and TATA box within this region. Insertion of the U6 promoters was confirmed via PCR and Sanger sequencing. Adapted plasmids are here on referred to as Llon1-pDCC6 and Ppap1-pDCC6. gRNA’s targeting genes of interest were inserted into the plasmid by Gibson Assembly to produce a panel of knockout plasmids (Supplementary Table S3). pDCC6 was a gift from Peter Duchek (Addgene plasmid # 59985; http://n2t.net/addgene:59985; RRID:Addgene_59985).

For piggyBac integration the Ubiq-Cas9.874W and pHome-T plasmids were selected. Ubiq-Cas9.874W contains a Ubi-63E promoter driving expression of hSpCas9 fused to GFP via a T2A self-cleaving peptide, and a fluorescent DsRed1 marker under control of the OpIE2 promoter. pHome-T expresses GFP under the synthetic promoter 3xP3 and RFP under control of the Ac5 promoter. Ubiq-Cas9.874W and pHome-T were gifts from Dr Roberto Galizi, (Keele University, UK) and Professor Tony Nolan (Liverpool School of Tropical Medicine, UK).

Two helper plasmids mhyPBase and ihyPBase, both hyperactive transposases, were delivered with piggybac plasmids containing a mammalian based transposase and Drosophila based hyperactive transposase respectively (Wright et al. 2013; Eckermann et al. 2018). Both hyperactive transposase plasmids were a gift from Dr Ernst Wimmer and Dr Hassan M.M. Ahmed (University of Gottingen, Germany).

### Rearing of sand flies

Sand fly colony was maintained as described previously (Volf and Volfova 2011). Briefly, *L. longipalpis* and *P. papatasi* adults were housed in cages, with access to 30% sugar solution at 26 - 28°C, on a 14 hr:10 hr light-dark cycle. Blood-fed females were separated into small cages for 5-6 days to defecate, gravid females were aspirated into plastic pots to lay eggs 6-10 days post blood-feeding and removed once oviposition had occurred. Larvae hatched 6-10 days later, with the larval period typically lasting ∼3 weeks followed by pupation for about 10 days Larval pots were placed within plastic boxes containing a base of moist sand (Lawyer et al. 2017).

### Microinjections

A 10μl stock injection mixture was prepared prior to each injection session and incubated at 37°C for 10-20 minutes, before being placed on ice for 20 minutes prior to injections. 2μl was loaded into the needle (G-1 borosilicate glass capillaries, outer diameter 1mm; inner diameter 0.6mm, Narishige, Japan). For PiggyBac transformation lines, mixtures consisted of a PiggyBac construct and hyPBase transposase helper plasmid in a 1:2 ratio. Injection mixtures for UbiqCas9.874W and pHome-T plasmids were prepared in the same manner (Supplementary Table S4).

For CRISPR knockouts, plasmids (pDCC6 variants) targeting the same gene were inoculated simultaneously. Three *Rudimentary*-targeting, and three *vestigial*-targeting constructs were combined in an even ratio to derive a final concentration of 317.58ng/μL. Separately, six olfactory-targeting plasmids were combined into a single injection mixture containing an even ratio of each to a final concentration of 316.68ng/μL. Additionally four ebony pDCC6 constructs were combined for a concentration of 278.48ng/μL (Supplementary Table S4).

Injections were performed using a manual Narishige IM-9B microinjector and Narishige MMO-4 micromanipulator (Narishige, Japan) coupled to an inverted microscope (Leica DMi8, Germany). Each egg was injected between 1/4 and 1/3 of the length of the egg from the posterior pole (Jeffries et al. 2018). Once injected, glass slides containing the eggs were transferred to a petri dish containing moist filter paper, and maintained at 25-28°C, 80% relative humidity. After ∼1-3 days eggs were brushed from the glass slides into oviposition pots.

### Identification of transgenic insects

A combination of methods were used to assess mutagenesis in transfected G0 survivors and subsequent G1 offspring. For emergent survivors transfected with plasmids inserting fluorescent cargo (PiggyBac plasmids) or targeting genes implicated in phenotypic change (CRISPR knockout plasmids) microscopy was performed. Secondly a PCR detection approach to identify mutagenesis was performed by T7 Endonuclease I Heteroduplex analysis. Thirdly a confirmatory *in silico* approach by Algorithmic deconvolution of Sanger sequence data was undertaken to predict indels.

### Fluorescent microscopy

Emergent transgenic insects were imaged by fluorescence stereo microscope (Leica M205 FA, Germany). L1-L4 larvae were imaged to identify fluorescence for GFP (485nm) and EGFP (488nm). DsRed1 and RFP markers imaged at 558nm and 587nm respectively. Positive outcomes for injection survivors of *UbiqCas9*.*874W* PiggyBac transfections is fluorescent mosaicism indicative of plasmid integration and expression (Opie2 promoter of DsRed), and green fluorescent mosaicism indicating expression of Cas9 (Cas9 tagged with GFP, separated by T2A ribosome skipping element).

### DNA extraction and PCR amplification

50% of emergent larvae were harvested for DNA extraction (DNeasy Blood and Tissue Kit, Qiagen, Germany). Sand flies without phenotypic changes by microscopy were pooled in batches of 10. DNA was extracted from individual sand flies if there was visual evidence of phenotypic change such as changes to wing morphology (genes targeting wing development or pigmentation). The remaining 50% were left to pupate until adults emerged for subsequent backcrossing. PCR amplification of extracted genomic DNA was performed in 20μl Phusion HF Master mix according to the manufacturer’s instructions (NEB, USA).

### Outcrossing/backcrossing

G0 male and female *L. longipalpis* injected with wing-targeting CRISPR constructs were sibling crossed, as were *L. longipalpis* individuals transfected with CRISPR-based olfactory constructs (Supplementary Table S3).

Additionally, two rounds of PiggyBac UbiqCas9.874W microinjections were performed. For the first round (*L. longipalpis and P. papatasi*), 50% of larvae were allowed to pupate and eclose (the remaining 50% of larvae had DNA extracted). G0 adults were separated by sex and out crossed with wildtype adults. Subsequent G1 larvae were microscopically examined for fluorescent phenotypes and DNA extracted for sequencing if they did not survive to G1 adults. In a second round of UbiqCas9.874W microinjections, all *L. longipalpi*s larvae were allowed to pupate and eclose, with adult survivors outcrossed with wildtype.

### Detecting mutagenesis by T7 Endonuclease I Heteroduplex assays

Mutagenesis was investigated using a T7 Endonuclease I Heteroduplex assay (T7EI, NEB, USA) according to the manufacturer’s instructions. Initially PCR amplifies the target region through successive rounds of denaturation and annealing with heteroduplexes forming between amplicons containing a modified locus and those with an unmodified locus. The subsequent pool of amplicons encompasses homoduplexes of unmodified DNA and heteroduplexes of mutated DNA. T7 Endonuclease recognizes mismatches (heteroduplexes) inducing double-strand breaks (DSB) with fragments detected by Gel electrophoresis followed by subsequent Densitometric analysis of amplicons in Image Lab software (Bio-Rad, USA). Fragments generated by T7EI cleavage, were defined using the Lane Profile tool (background excluded) and the Analysis Table function to generate the Adjusted volume (background excluded volume) and the Band Percent (percent of the adjusted volume of the selected band, relative to the other selected bands in the lane (% density of the sum of the Adjusted volume).

### ICE analysis, algorithmic deconvolution analysis of sequence data to detect mutagenesis

Sanger sequencing chromatograms were analyzed using ICE (Inference of CRISPR Edits), an algorithmic trace-deconvolution method that estimates CRISPR editing outcomes from mixed amplicon populations by comparing edited samples to unedited control traces. ICE analysis was performed *in silico* using the Synthego Performance Analysis (ICE Analysis; v3.0, 2019). Briefly, Sanger files were uploaded together with the relevant guide RNA sequences. The software inferred editing efficiency and the spectrum of indels for knockout experiments. Workflow was used to assess intended sequence insertion/replacement events (up to ∼200 bp).

## Results

### Microinjection Efficiency and Survival

A total of 7,623 sand fly embryos (*L. longipalpis* and *P. papatasi*) were injected with PiggyBac transposon constructs to integrate GPF or DsRed1 markers or CRISPR plasmids (Table 1) targeting wing-development, pigmentation, immunity, or olfactory genes (Table 2). Survival rates were comparable to those previously published, with adult survival as a percentage of eggs injected from 1.04 - 4.81%.

**Table 1.** Survival for *L. longipalpis* and *P. papatasi* eggs injected with multiple plasmids. Counting larvae is difficult due to substrate and can result in undercounting in comparison to emergent sand flies, occasionally inferring adult survival as % of larvae being greater than 100% (indicated by *†)*.

| *Species* | *Plasmid(s)* | *Injected*<br>*eggs* | *Larval*<br>*survival*<br>*(%)* | *Adult*<br>*survival*<br>*Male* | *Adult*<br>*survival*<br>*Female* | *Adult*<br>*survival*<br>*as% of*<br>*larvae* | *Adult*<br>*survival*<br>*as% of*<br>*eggs*<br>*injected* |
| --- | --- | --- | --- | --- | --- | --- | --- |
| *L.*<br>*longipalpis* | UbiqCas9.874W<br>+ IhyPBase | 2058 | 75<br>(**3.64**) | 27 | 15 | 56.0 | 2.04 |
| *P. papatasi* | UbiqCas9.874W<br>+ IhyPBase | 1827 | 44<br>(**2.41**) | 29 | 13 | 95.45 | 2.30 |
| *L.*<br>*longipalpis* | pHome +<br>IhyPBase | 749 | 57<br>(**7.61**) | 15 | 21 | 63.16 | 4.81 |
| *P. papatasi* | pHome +<br>IhyPBase | 800 | 42<br>(**5.25**) | 12 | 9 | 50.0 | 2.63 |
| *P. papatasi* | pDCC6:<br>Ebony1,<br>Ebony2,<br>Ebony3, Ebony4 | 1034 | 8 (**0.77**) | 6 | 6 | 150† | 1.16 |
| *L.*<br>*longipalpis* | pDCC6:<br>R1, R2, R3, V1,<br>V2, V3 | 579 | 6 (**1.04**) | 3 | 3 | 100 | 1.04 |
| *L.*<br>*longipalpis* | pDCC6:<br>Gr2-1,<br>Gr2-2,<br>Ir8a-1,<br>Ir8a-2,<br>Orco 12,<br>Orco 34 | 576 | 10<br>(**1.74**) | 9 | 2 | 110† | 1.91 |

**Table 2.** Summary of evidence for CRISPR mutagenesis across gene targets (*Lutzomyia longipalpis* and *Phlebotomus papatasi*). Evidence types encompass phenotypic mutantagenesis, detection by T7 Endonuclease I heteroduplex cleavage assays, with subsequent densitometric quantification and *in silico* sequence analysis by ICE algorithmic indel prediction. Mutations were detected across both G0 and G1 generations depending on the target and method. Rudimentary, *Gr2*, and *Orco* demonstrate robust multi-method support indicative of mutagenesis.

| *Gene Target* | *Function / Class* | *Cut Sites / Constructs* | *Evidence of Mutagenesis* | *Generations* | *Comment* |
| --- | --- | --- | --- | --- | --- |
| *Rudimentary (r)* | Wing development | R2, R3 | ICE indels; T7E1; Wing phenotype mutants | G0, G1 | Strong evidence across assays |
| *Vestigial (vg)* | Wing development | V1, V2, V3 | T7E1 (V1,V2); limited ICE | G1 | Sequencing limited V3 detection |
| *Ebony (e)* | Pigmentation | Ebony constructs | Darkened G0 pupae (phenotype) | G0 | Phenotypic confirmation only |
| *Gr2* | Olfaction | Gr2-1, Gr2-2 | ICE indels; T7E1 | G0, G1 | Strong olfactory evidence |
| *Ir8a* | Olfaction | Ir8a-1, Ir8a-2 | ICE indels; T7E1 | G0, G1 | Moderate but consistent |
| *Orco* | Olfaction (co-receptor) | Orco-1, Orco-2 | ICE indels; T7E1; High-confidence KO (G1-MA) | G0, G1 | Most robust knockout evidence |

### Evidence of PiggyBac Integration of Cas9 sequences

Microinjection with two different PiggyBac plasmids pHOME-T and UbiqCas9.874W into *L. longipalpis* and *P. papatasi* embryos resulted in successful genomic integration, detectable at the molecular level (via PCR detection and Sanger sequencing) in both the G_0_ and G_1_ generations, despite the absence of 3xP3-driven GFP or Actin5C driven RFP fluorescent marker expression in emergent individuals. In more detail, for pHOME-T, molecular screening identified genomic integration in G_0_ pooled larval samples of both vector species. Germline transmission was also confirmed through analysis of G_1_ progeny by Sanger sequencing. In *L. longipalpis*, 9 of 60 pooled G_1_ larval samples were GFP-positive by PCR, with 7 of these 9 further validated by Sanger sequencing. In *P. papatasi*, 7 of 14 pooled G_1_ larval samples were GFP-positive, with all seven confirmed by sequencing. Together, these results demonstrate both integration and inheritance of the pHOME-T cargo.

For UbiqCas9.874W, PCR and sequencing confirmed integration of Cas9, GFP, or DsRed1 sequences in G_0_ larvae and adults of both sand fly species. Germline transmission was observed in *L. longipalpis*, where 33 of 52 G_1_ pooled larval samples were PCR-positive for Cas9, and 15 of these were confirmed by sequencing (Supplementary Figure S5). For G0 *P. papatasi* injected with UbiqCas9.874W and IhyPBase (1,827 injected), two pooled samples of 10 larvae and two adult samples were PCR positive for Cas9, and two single adult samples were also positive for Cas9 via PCR and sequencing. No G_1_ positives were detected for *P. papatasi*, in line with limited G_0_ survival in this species. No DsRed1 or EGFP expression was detectable.

These data provide the first demonstration that PiggyBac cargo can integrate into sand fly genomes of both species tested and that exogenous cargo can be inherited through the germline. The current work validates these constructs for genetic manipulation in both *L. longipalpis* and *P. papatasi*.

### Evidence for CRISPR-Cas9 Induced mutagenesis

#### Phenotypic Mutagenesis

Evidence of mutagenesis affecting wing development (*L. longipalpis*) in three G1 individuals exhibited observable wing deformities, namely 1 with unilateral wing loss and the remaining two with pointed wings consistent with expected CRISPR disruptions (Figure 1). Importantly, phenotypes were not only observed visually but were also supported by DNA-level evidence of CRISPR mutagenesis, although the strength of molecular confirmation varied among individuals (Table 2). The individual with unilateral wing loss presented T7E1 heteroduplex cleavage (R2/R3 target) with subsequent support from ICE analysis (R3 target) comprising strong evidence of CRISPR targeted mutagenesis. Two additional flies (pointed wings) also demonstrated T7E1 mismatch evidence, but without supporting ICE analysis, due to sequence quality, and so a more moderate evidence level. Overall, phenotypic mutants had molecular support from at least one assay.

**Figure 1.**
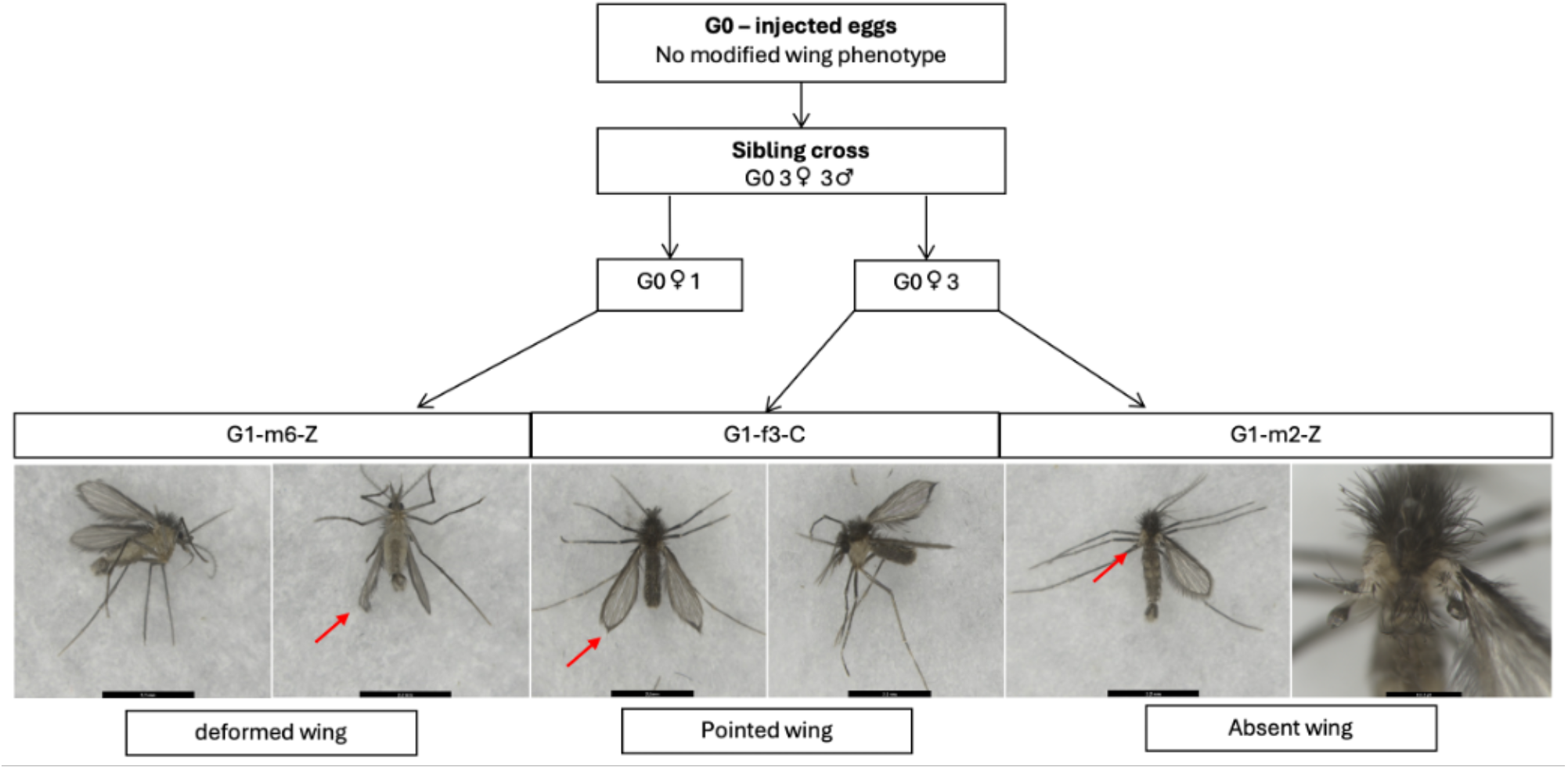
G1 microinjection survivors with atypical wing phenotypes. Phenotypic characteristics of G1 male individual (sample G1-m6-Z) presenting a deformed wing phenotype, G1 female individual (sample G1-f3-C) presenting pointed wing tip phenotype, and G1 male (sample G1-m2-Z) undeveloped left wing. Scale bars 2.2mm.

### Molecular Evidence of CRISPR Mutagenesis

#### T7 Endonuclease I Heteroduplex Assays Reveal Evidence of CRISPR Editing

Sanger sequencing alone is not reliable to identify CRISPR-Cas9 modification where the dominant sequence can mask minority modified alleles. Therefore, to detect minority insertions and deletions generated by CRISPR editing, T7 endonuclease I (T7EI) assays were performed on PCR amplicons spanning targeted cut sites in *L. longipalpis* G_0_ and G_1_ individuals.

For wing-targeting constructs, amplicons were successfully obtained for all G_1_ samples at the R3 (534 bp), V1 (577 bp), and V2 (567 bp) loci. Heteroduplex cleavage was observed in six R3 samples (R3C*, R3M*, R3N*, R3U*, R3V*, R3Z*) and in six V1 and nine V2 samples (Figure 2), each producing digestion fragments between 200–400 bp consistent with CRISPR-derived mismatches.

**Figure 2.**
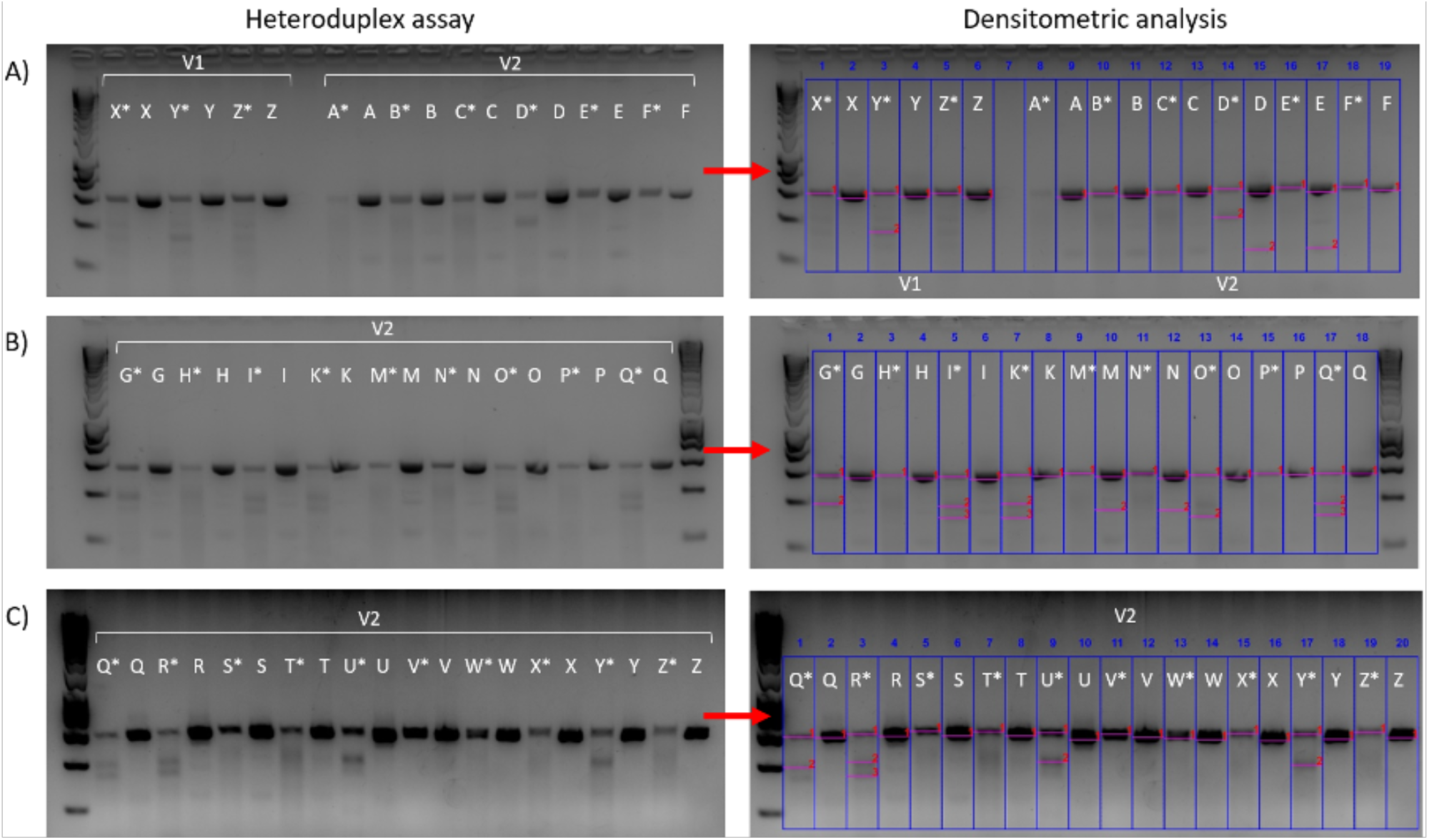
*L. longipalpis* T7 Endonuclease I Heteroduplex assay and densitometric analysis (G1samples targeting V1 and V2 gRNAs regions). A) Samples V1X-V1Z followed by V2A-V2F, B) samples V2G-V2Q, and C) samples V2Q-V2Z. Asterisks (*) indicates PCR product and subsequent digestion with endonuclease. Lanes without asterisk are controls untreated with Endonuclease. For Densitometric analysis (an estimate of the proportion of modified alleles) blue boxes indicate boundaries of each lane, and pink lines indicate amplicons assessed within each lane.

In olfactory-gene targeted flies, T7EI assays similarly provided evidence of mutagenesis. Among G_0_ individuals, six Gr2 and four Ir8a amplicons showed digestion consistent with heteroduplex formation, while in G_1_ samples, T7EI activity was detected across *Gr2, Ir8a*, and *Orco* loci. Notably, nine male and nine female G_1_ samples exhibited heteroduplex digestion for *Orco*, with fragments between 200– 600 bp.

#### Densitometric Quantification Indicates Low-to-Moderate Modification Levels

To estimate the proportion of modified alleles within each amplicon pool, densitometric analysis was applied to T7EI digests. For wing-gene targets, modification rates at R3 ranged from 0.36–6.13%, whereas V1 and V2 targets showed broader editing ranges, from 2.98–18.51% and 0.64–15.06%, respectively. Background digestion in controls remained consistently low (0.05–4.43%) and is highly suggestive that elevated cleavage intensities in experimental samples reflected genuine CRISPR editing rather than nonspecific nuclease activity.

Olfactory-gene knockouts showed similarly variable evidence of modification. In G_0_ flies, densitometry estimated 0.68–7.52% modification at Gr2 and 0.48–3.91% at Ir8a, while G_1_ samples demonstrated wider ranges. Gr2 modification ranged from 0.19% to 57.64%, and Orco modification ranged from 0.44–9.95%, exceeding background digestion levels in named samples.

#### ICE Sequence Deconvolution Independently Supports CRISPR Editing

Inference of CRISPR Edits (ICE) analysis provided an orthogonal method to detect indels directly from Sanger sequencing traces.

For wing-targeting constructs, ICE predicted indels at the R2 cut site in 16 of 26 G_1_ individuals, including those displaying wing deformities, and in two individuals at R3.

Olfactory gene analyses further support editing events. ICE detected indels in 8 of 11 G_0_ individuals and in 17 of 34 G_1_ individuals at Gr2, with 10 of these 17 showing matching T7EI cleavage, representing strong evidence of true gene editing. For Ir8a, 17 G_1_ samples showed ICE predicted indels, and three were corroborated by T7EI. For Orco, 25 G_1_ individuals carried ICE predicted indels, including one (G1-MA) with exceptionally strong support (R^2^ = 0.99, KO-score = 1) and concordant T7EI digestion.

Together, the T7EI heteroduplex assays, densitometric quantification, and ICE-based sequence analysis provide strong molecular evidence of CRISPR-mediated mutagenesis in both G_0_ and G_1_ *L. longipalpis*. Although individual assays varied in sensitivity and specificity, the combined dataset reveals consistent patterns of indel formation across multiple target genes, with editing detected at wing-development loci (R3, V1, V2) and olfactory genes (Gr2, Ir8a, Orco). Crucially, numerous individuals exhibited concordance between two or more methods, including 18 instances where T7EI and ICE jointly identified indels, providing robust confirmation that CRISPR-Cas9 delivered via plasmid microinjection successfully induced heritable genome editing in sand flies.

### Summary of Editing Outcomes

PiggyBac-Cas9 delivery using two different plasmid constructs (UbiqCas9.874W and pHOME-T) successfully produced germline-transmitting lines. The former verified Cas9 integration in G0 and G1 individuals, and the latter GFP expression in (G1), confirmed by Sanger sequencing. Wing-development genes showed both genotypic mutations and heritable phenotypic defects. For CRISPR - induced mutagenesis, olfactory genes (*Gr2, Ir8a, Orco*) showed clear evidence of CRISPR editing in G0 and G1 individuals (Table 2). Overall, these results demonstrate the first successful *in vivo* genetic modification of *L. longipalpis* and *P. papatasi* using PiggyBac-mediated integration of exogenous DNA, as well as the first reported CRISPR-Cas9 knockout mutagenesis in *L. longipalpis*.

## Discussion

Functional genetic tools for phlebotomine sand flies significantly lag behind those available for other insects of medical importance, particularly mosquito vectors, which has hampered progress toward development of novel genetic vector control strategies for leishmaniasis. This is despite the continued epidemiological impact of the disease.

In recent years a plethora of publications have described CRISPR-Cas9 gene editing applied to insects of medical, veterinary and agricultural importance, stemming from initial demonstration in *Drosophila melanogaster* (Gantz and Akbari 2018; Gratz et al. 2014; Gokcezade et al. 2014). Notable examples include Anopheline vectors of malaria, *Aedes* vectors of dengue and Zika viruses, and major crop pests *Plutella xylostella* and *Ceratitis capitata*, amongst many others (Gantz and Akbari 2018).

Early publications often described species-specific protocols exploring functional genomics in the context of gene knockouts (Kistler et al. 2015; Basu et al. 2015; Gantz et al. 2015; Hammond et al. 2016). More recently, refined homology-directed repair approaches have been developed in mosquito species to facilitate gene drives, enabling the spread of introduced heritable traits through populations at levels exceeding those achievable by Mendelian inheritance. Broadly, there are two main implementations of gene-drives applied to insects of medical importance, both interrupting disease transmission. The first, population suppression strategies, attempt to reduce the vector populations below that required to sustain pathogen transmission (Hammond et al. 2016; Simoni et al. 2020b)(Simoni et al. 2020a; Bernardini et al. 2018). Typically, the strategy introduces lethal or deleterious self-propagating traits affecting overall reproductive fitness leading to drastic reduction in local populations. An alternative approach, referred to as population replacement or modification, is transmission blocking, for example expression of antiparasite peptides that target vector genes implicated in establishing or transmitting infection to derive pathogen-resistant vectors (Gantz et al. 2015; Williams et al. 2020). Each approach has its benefits and drawbacks. Technical challenges remain (Hammond, Karlsson, et al. 2021; Garrood et al. 2021)(Beaghton et al. 2019); however, both approaches offer exciting potential to deliver tangible results and show great promise in laboratory settings as well as large-scale caged studies (Bier 2022; Hammond, Pollegioni, et al. 2021).

PiggyBac is the most widely used transposon system for genetic manipulations in insects, successfully applied to many different mosquito vectors to express fluorescent markers and anti-parasite peptides (Nolan et al. 2002; Grossman et al. 2000)(Ito et al. 2002; Nolan et al. 2002)(Perera et al. 2002)), in *Aedes aegypti* (Kokoza, Ahmed, Wimmer, & Raikhel, 2001), *Ae. albopictus* (Labbé, Nimmo, & Alphey, 2010), and *Ae. fluviatilis* (Rodrigues, Oliveira, Rocha, & Moreira, 2006). PiggyBac plasmids can carry CRISPR-Cas components as cargo for insertion into genomes (Li et al. 2017). Importantly, the technique has not been demonstrated in sand flies which is a significant omission for such a potentially useful tool.

In the current study, we establish a robust *in vivo* framework for genetic modification of *L. longipalpis* and *P. papatasi*. We provide the first evidence of PiggyBac-mediated transgenesis in *L. longipalpis* and *P. papatasi* and also CRISPR-Cas9-based targeted mutagenesis in *L. Longipalpis*, the major vector of visceral leishmaniasis in the Americas.

For PiggyBac constructs incorporating a ubiquitin-driven Cas9 cassette, Cas9 and GFP sequences were detected in both G0 and G1 individuals, together indicating successful germline transmission and establishing PiggyBac as a viable transformation system in sand flies. This finding addresses an important technical limitation highlighted by previous unsuccessful attempts described in the literature for *L. longipalpis* (Martin-Martin et al. 2018; Jeffries et al. 2018; Louradour et al. 2019). Building on this, we describe a functional approach to support future gene editing studies.

In parallel, the results provide multiple lines of evidence supporting CRISPR-Cas9-mediated mutagenesis that target both phenotypic marker genes and olfactory genes.

Molecular analyses by Sanger sequencing suggest that mosaicism is a dominant feature of CRISPR editing in sand flies, as expected given plasmid-based delivery during early embryogenesis. Overall, standard Sanger sequencing alone proved insufficient to reliably detect low-frequency edits. However, the combined application of T7 endonuclease I assays, densitometric analysis, and in silico ICE deconvolution applied to Sanger sequences provided robust and reproducible evidence of targeted mutagenesis by application of CRISPR based approaches. Considering that rapid pre-cellular nuclear divisions occur shortly after oviposition in dipterans, reducing the 4-hour time window between collecting eggs and injecting them, if logistically possible, may help reducing mosaicism (Jeffries et al. 2018).

Targeting olfactory genes (*Gr2, Ir8a*, and *Orco*) implicated in chemosensory function, host-seeking behaviour, and the maintenance of vectorial capacity across Diptera, confirms that key behavioural pathways are, in principle, genetically accessible in sand flies. These findings provide a foundation for investigating the modulation of host seeking behaviour and exploring strategies to interrupt parasite transmission.

Low recovery of phenotypically visible mutants likely reflects a combination of mosaicism, developmental lethality or impaired adult fitness, together with the intrinsic challenges of sand fly embryo microinjection. Although phenotypic mutants were recovered at low frequency, observed wing malformations and pigmentation changes were consistent with predicted orthologous knockouts in mosquitoes. In this context, edits supported by molecular evidence were biologically consistent with disruption of vestigial (*vg*) and rudimentary (*r*).

In summary, this work establishes the first integrated approach for sand fly genetic modification, bridging bioinformatic target identification, the successful application of molecular tools incorporating piggyBac and CRISPR based approaches, and subsequent *in vivo* validation. The recent availability of improved assemblies for *P. papatasi* and *L. longipalpis* (Huang et al. 2024) will facilitate future targeting efficiency. Together, these results represent a critical step toward the application of advanced genetic control strategies for leishmaniasis vectors and will contribute to ongoing behavioural, ecological, and importantly, gene-drive development to facilitate vector control.

## Acknowledgments

We are grateful to Hassan M. M. Ahmed, Ernst Wimmer, Tony Nolan and Roberto Galizi for the donation of plasmids.

## Author contributions

M.Y., conceptualised the study; R.T.M.E., L.B.G., B.V.K., and E.L.T. performed all experiments; M.Y., R.T.M.E., L.B.G., B.V.K., and E.L.T. performed data analysis; M.Y., and R.T.M.E., wrote the first draft of the manuscript; T. B. reviewed and edited an advanced draft of the manuscript; M.Y. T.W., M.E.R., and P.V provided resources and supervision; All authors contributed to analysing and compiling the data, writing and approving the final manuscript.

