## Supplementary material for "CRISPR-Cas9 and PiggyBac Mediated Genetic Modification of Sand Fly Vectors Targeting Olfactory and Non-Lethal Phenotypic Genes": Table S1

**Table S1.** Genetic loci identified for CRISPR-Cas9 gene editing with associated orthologues of *L. longipalpis* and *P. papatasi*.

| ***Gene*** | ***Phenotype on knockout/Kairomone response*** | ***L. longipalpis orthologue*** | ***P. papatasi orthologue*** |
| --- | --- | --- | --- |
| ***ebony (e)*** | Suppresses melanin formation in specific cuticle cells. | LLOJ008326 | PPAI005863 |
| ***rudimentary (r)*** | wing malformations and pyrimidine auxotrophy | LLOJ009278 | N/A |
| ***vestigial (vg)*** | encodes a wing/haltere identity selector gene | LLOJ009695 | PPAI008343 |
| ***Orco*** | Co-receptor | LLOJ003114 | PPAI006929 |
| ***Gr2*** | CO_2_ | LLOJ010835 | PPAI013119 |
| ***Ir8a*** | Acids and amine (co-receptor) | LLOJ003646 | PPAI001607 |
