## Supplementary material for "CRISPR-Cas9 and PiggyBac Mediated Genetic Modification of Sand Fly Vectors Targeting Olfactory and Non-Lethal Phenotypic Genes": Figure S2

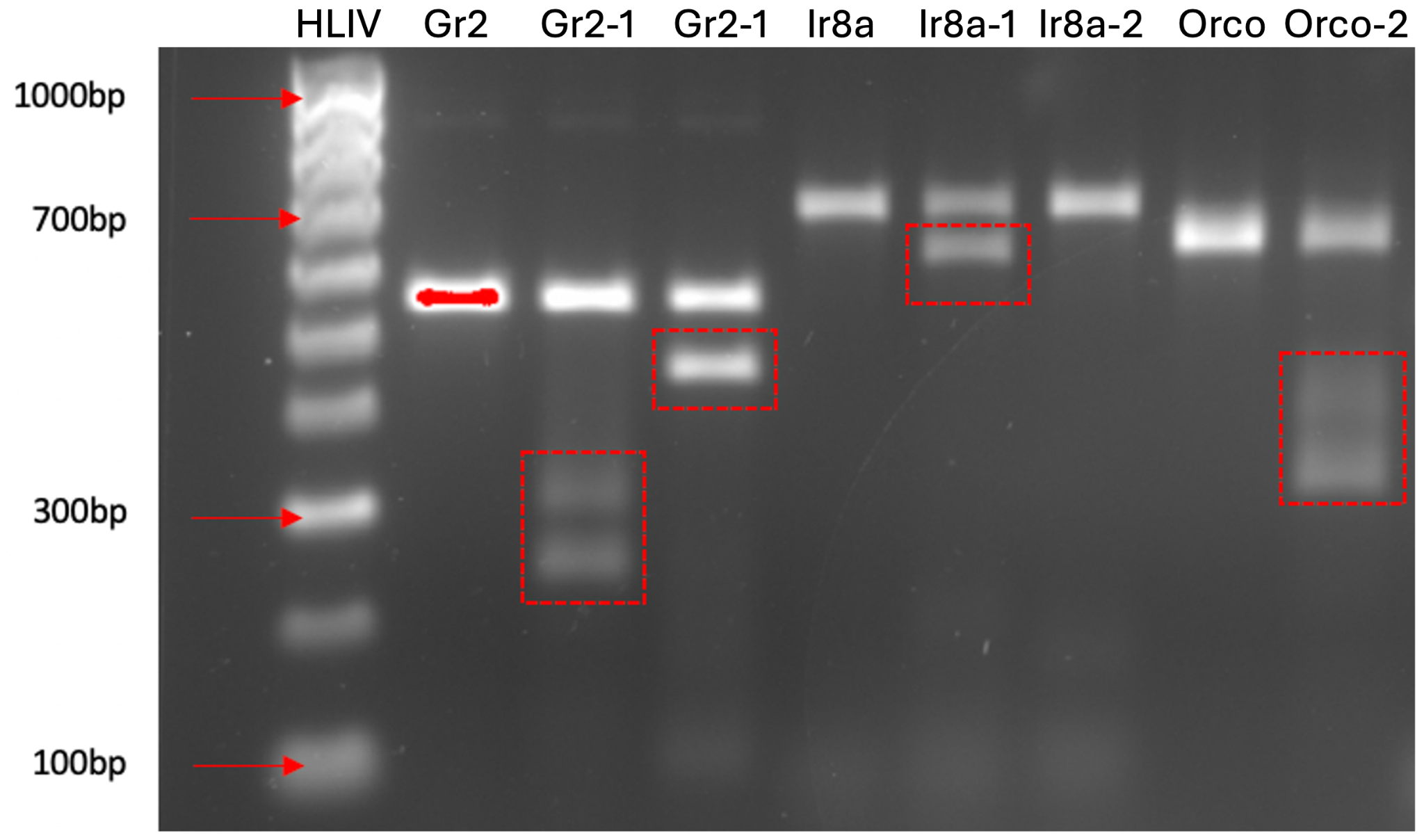

**Figure S2**. *In vitro* Endonuclease Cleavage Assay of olfactory targets s*gRNAs* *Gr2-1*, *GR2-2*, *Ir8a-1*, *IR8-2* and *Orco*. Controls (no sgRNAs) loaded next to lanes with digested samples sgRNAs: Gr2 control (gr2) (*lane 2*), Ir8a control (ir8a) (*lane 5*); Orco control (orco) (*lane 8*).
