## Supplementary material for "CRISPR-Cas9 and PiggyBac Mediated Genetic Modification of Sand Fly Vectors Targeting Olfactory and Non-Lethal Phenotypic Genes": Table S3

**Table S3.** pDCC6 CRISPR knockout constructs targeting olfactory, wing development, and pigmentation genes in *L. longipalpis* and *P. papatasi.* gRNA (green) with overhangs (black) for Gibson assembly into the pDCC6 backbone. AAAC and AATT are overhangs for *L. longipalpis*, and AATT and AAAC are overhangs for *P. papatasi.*

| ***Species*** | ***Gene*** | ***gRNA sequence*** | ***Construct name*** |
| --- | --- | --- | --- |
| ***L. longipalpis*** | Gr2 | AAAC ATAAAGAGGCAATACGTGG C | Gr2-1-pDCC6 |
|  |  | AATT G GCTGTGTACAAGACAATGTGGGG | Gr2-2-pDCC6 |
|  | Orco | AATT G TGTGAGATACATGACCAACAAGG | Orco-1-pDCC6 |
|  |  | AATT G TACAGCAATCAAGTATTGGGTGG | Orco-2-pDCC6 |
|  | Ir8a | AAAC AGCAATAGCGAGACCTGCGAGGG C | Ir8a-1-pDCC6 |
|  |  | AATT G CCCTGTCGGGATTTTACACTCAA | Ir8a-2-pDCC6 |
|  | Rudimentary | AATT AGCATTGGAAGACACAGGGT | R1-pDCC6 |
|  |  | AATT TTGTAGCCCATCTTCACGA | R2-pDCC6 |
|  |  | AATT GGAACTATGGCATTCCGTG | R3-pDCC6 |
|  | Vestigial | AATT TCATCATTACGGTTCCTACG | V1-pDCC6 |
|  |  | AATT GGAAAATTTCTCGCCGACAT | V2-pDCC6 |
|  |  | AATT TCGCGGACACGTATTGTGCT | V3-pDCC6 |
| ***P. papatasi*** | Ebony | ATTT TCGCATTCAGCACATCCTTG | Ebony1-pDCC6 |
|  |  | ATTT AAAGTGCATGGTAATCAGGA | Ebony2-pDCC6 |
|  |  | ATTT CCTGGCCATATGGAAATGTG | Ebony3-pDCC6 |
|  |  | ATTT TCGGACAACTTGATAGCCAC | Ebony4-pDCC6 |
