## Supplementary material for "CRISPR-Cas9 and PiggyBac Mediated Genetic Modification of Sand Fly Vectors Targeting Olfactory and Non-Lethal Phenotypic Genes": Table S4

**Table S4**. Injection mixtures for microinjection of *L. longipalpis* *and P. papatasi* eggs.

| ***Plasmid*** | ***Actual conc. (ng/µl)*** | ***Final Conc. (ng/µl)*** | ***Mixture conc.(ng/µl)*** | ***Plasmid Ratio*** |
| --- | --- | --- | --- | --- |
| ***UbiqCas9.874W*** | 82.9 | 19.98 | 61.41 | 1;2 |
| ***IhyPBase*** | 414.3 | 41.43 |  |  |
| ***UbiqCas9.874W*** | 98.5 | 20.00 | 59.86 | 1;2 |
| ***IhyPBase*** | 538.7 | 39.86 |  |  |
| ***UbiqCas9.874W*** | 98.5 | 39.99 | 120.26 | 1;2 |
| ***IhyPBase*** | 538.7 | 80.27 |  |  |
| ***UbiqCas9.874w*** | 80.7 | 62.10 | 186.29 | 1;2 |
| ***IhyPBase*** | 538.7 | 124.19 |  |  |
| ***pHOME*** | 449.5 | 20.23 | 61.66 | 1;2 |
| ***IhyPBase*** | 414.3 | 41.43 |  |  |
| ***pHOME*** | 449.5 | 40.46 | 123.32 | 1;2 |
| ***IhyPBase*** | 414.3 | 82.86 |  |  |
| ***R1*** | 299.1 | 52.93 | 317.58 | 1;1;1;1;1;1 |
| ***R2*** | 335.6 | 52.93 |  |  |
| ***R3*** | 370.6 | 52.93 |  |  |
| ***V1*** | 324.7 | 52.93 |  |  |
| ***V2*** | 405.3 | 52.93 |  |  |
| ***V3*** | 231.2 | 52.93 |  |  |
| ***Gr2-1*** | 335.5 | 52.78 | 316.68 | 1;1;1;1;1;1 |
| ***Gr2-2*** | 342.3 | 52.78 |  |  |
| ***Ir8a-1*** | 278 | 52.78 |  |  |
| ***Ir8a-2*** | 345.6 | 52.78 |  |  |
| ***Orco 12*** | 265.2 | 52.78 |  |  |
| ***Orco 34*** | 359.5 | 52.78 |  |  |
| ***Ebony gRNA 1*** | 314 | 69.62 | 278.48 | 1;1;1;1 |
| ***Ebony gRNA 2*** | 164 | 69.62 |  |  |
| ***Ebony gRNA 3*** | 473 | 69.62 |  |  |
| ***Ebony gRNA 4*** | 337 | 69.62 |  |  |
