## Supplementary material for "CRISPR-Cas9 and PiggyBac Mediated Genetic Modification of Sand Fly Vectors Targeting Olfactory and Non-Lethal Phenotypic Genes": Figure S5

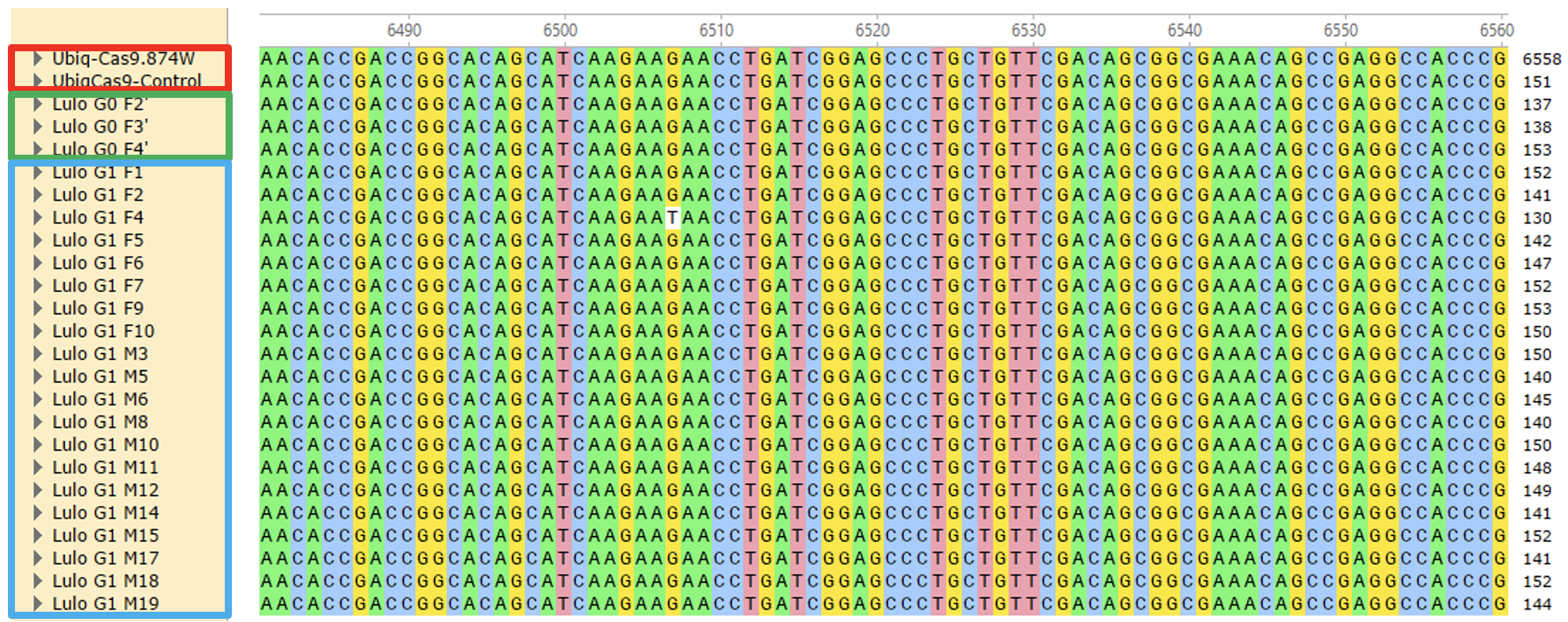

**Figure S5.** Inserted Cas9 cargo sequence alignment of G0 and G1 *L. longipalpis* samples injected with PiggyBac UbiqCas9.874W. G0 samples (green box) and G1 samples (blue box) derived from G0 and wildtype crosses. Sample sequences were aligned with the Cas9 sequence derived from the UbiqCas9.874 W plasmid map, against Cas9 sequence amplified from the plasmid (control, red box).
